# SegMantX: a novel tool for detecting DNA duplications uncovers prevalent duplications in plasmids

**DOI:** 10.1101/2025.03.14.643293

**Authors:** Dustin M. Hanke, Tal Dagan

## Abstract

Segmental duplications play an important role in genome evolution via their contribution to copy-number variation, gene-family diversification and the emergence of novel functions. The detection of segmental duplications is challenging due to heterogeneous amelioration of sequence similarity among duplicates, which hinders the reconstruction of continuous sequence alignment. Here we introduce SegMantX, a novel approach for the identifcation of diverged segmental duplications using local alignment chaining. In this approach, local alignments resulting from a preliminary sequence similarity search (e.g., BLASTn) are chained into continuous segments. Evaluating the performance of SegMantX simulated sequences shows that the tool can detect diverged duplications beyond the sensitivity limits of standard alignment-based methods. Applying SegMantX to 6,784 enterobacterial plasmids, we find that 74% plasmids contain duplicated regions, most of which correspond to duplicated mobile genetic elements (MGEs; e.g., transposons and insertion sequences). Furthermore, we demonstrate the applicability of SegMantX for the identification of diverged gene transfers between replicons, and plasmid hybridization events. Our findings highlight MGEs as drivers of segmental duplications in plasmid evolution, leading to the amplification of their cargo genes, including antibiotic resistance genes. SegMantX provides a powerful framework for reconstructing diverged segmental duplications and other alignment problems.

## Introduction

Gene and whole genome duplications (WGDs) are fundamental mechanisms of genome evolution due to their contribution to gene family diversification (Ohno 1970; Van De Peer, Fawcett, et al. 2009). Duplications occur across multiple scales, including domains (or exons), genes, segmental duplications spanning multiple genes, and whole genome duplications (WGDs). The presence of duplicated genes may increase the gene product dosage. Duplicated gene copies may evolve distinct functions via neo- or sub-functionalization, or become non-functional copies (pseudogenes). In eukaryotes, gene duplications and WGDs, which are mainly driven by recombination, have been pivotal in speciation processes (Lynch and Conery 2000; Van De Peer, Maere, et al. 2009). For example, *Saccharomyces cerevisiae* emerged from a WGD event, followed by differential gene loss in duplicated gene copies, which contributed to the speciation of yeast lineages descending from a shared common ancestor (Wolfe and Shields 1997; Kellis et al. 2004; Scannell et al. 2006). In prokaryotes, the diversification of gene families is rather driven by horizontal gene transfer while duplication events are instead considered rare (Mahillon and Chandler 1998; Ochman et al. 2000; Lerat et al. 2005; Treangen and Rocha 2011; Tria and Martin 2021). Mobile genetic elements are often associated with gene duplications in prokaryotes, e.g., as shown for the evolution of plasmids in the nitrogen-fixing *Burkholderia vietnamiensis* (Maida et al. 2014).

Mobile genetic elements contribute also to the emergence of transient gene duplications in prokaryotes that is termed gene amplification. For example, tandem insertions of the TEM-1 beta-lactamase gene in an *E. coli* plasmid led to an increased gene dosage, thereby enhancing the strain resistance to ampicillin (Dhar et al. 2014). Transient amplification of resistance genes can lead to antibiotic heteroresistance, where genetically identical bacterial populations contain subpopulations with varying antibiotic resistance level (Nicoloff et al. 2024). MGEs thus play a major role in gene amplification and duplication in plasmid evolution, nonetheless, many MGE-mediated gene duplications remain as non-functional gene copies (Hanke et al. 2024). Large-scale duplications such as plasmid multimerization is analogous to whole-genome duplications. Their maintenance depends on selective pressure for the plasmid presence, as demonstrated in experimental *E. coli* evolution (Wein et al. 2020; Abe et al. 2021).

Methods for detection of DNA duplication are either based on copy number variation in next-generation sequencing data (e.g., DuplicationDetector, Djedatin et al. 2017) or sequence similarity in comparative genomics studies (e.g., BLAST and DIAMOND, Camacho et al. 2009; Buchfink et al. 2015). Advanced approaches further assess gene paralogy and orthology (e.g., OrthoMCL, OrthoFinder, or TreeFam, Li et al. 2003; Li 2006; Emms and Kelly 2019) or integrate functional annotation of the duplicated genes (e.g., HSDFinder, Zhang et al. 2021). Considering gene order conservation is furthermore useful for the identification of segmental duplications and WGDs (e.g., DAGChainer and MCScanX-transposed, Haas et al. 2004; Wang et al. 2013). Most of these tools were developed for the detection of duplicated protein-coding genes with a high sequence similarity, hence their applicability for detecting diverged duplications is limited. The use of gene-based detection is furthermore limiting the ability to detect DNA duplications in the presence of genomic rearrangements, horizontal gene transfer, and degraded sequence similarity among duplicated copies.

Here we present a novel method for the detection of DNA duplications – SegMantX – using local alignment chaining, and demonstrate its application to enterobacterial plasmids. The algorithm’s performance was validated with simulated data, demonstrating its effectiveness in reconstructing diverged duplications. Analyzing DNA duplications in plasmid genomes, we uncover the prevalence and characteristics of duplication events in plasmid evolution.

## Results

### Detection of duplicated genomic regions with local alignment is limited due to sequence divergence

For the development of our method and the demonstration of its performance we utilized a dataset of 6,784 plasmids reported in 2,441 complete enterobacterial genomes, including the genera *Klebsiella, Escherichia*, and *Salmonella* (*KES*) (Wang and Dagan 2024). Putative duplicated regions in plasmid genomes were preliminarily identified by sequence similarity search of the plasmid sequence against itself with BLASTn (Camacho et al. 2009). Using this approach, we detected putative duplication events in 5,043 (74%) plasmid genomes.

A close examination of plasmids where putative duplications were detected, often revealed the presence of consecutive local similarity hits including syntenic gene order. Example is a 187,669 kbp large plasmid in an *E. coli* strain isolated from waste water (Fig. 1A). Since the plasmid was assembled as a single contig, we reasoned that this sequence similarity pattern was likely the result of a single duplication event followed by sequence divergence. The application of our algorithm to the plasmid resulted in chaining of the 13 BLAST hits into a single segment with a global sequence similarity of 87% identical amino acids (Fig. 1B). The ∼60 kbp segment comprises 74 genes of plasmid core functions, primarily related to plasmid transfer and replication (Fig. 1C, Supplementary Table 1). The syntenic gene arrangement in the duplicated region points towards a large segmental duplication followed by sequence divergence. The genes in this segmental duplication were mostly retained with high sequence similarity between the duplicated (Fig. 1C). The recovered segment corresponds conceptually to a self-sequence alignment of the plasmid genome; however, it could have not been fully reconstructed by standard local (or global) alignment methods due to the heterogeneity of sequence divergence post-duplication event.

**Figure 1.**
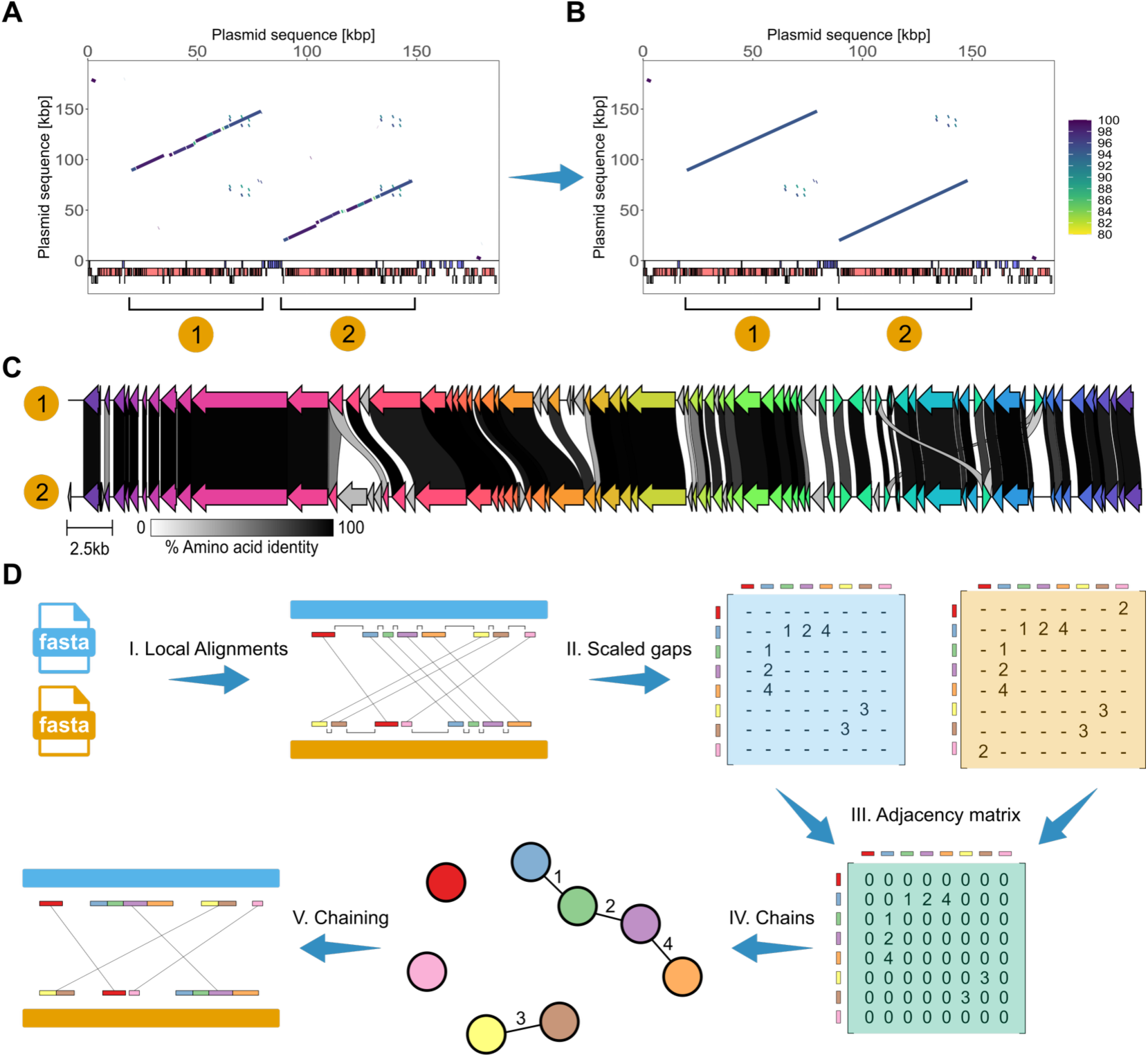
Detection of diverged segmental gene duplications via chaining of local similarity alignments. **(A)** Result of sequence similarity alignment of plasmid sequence used both as query and subject with BLASTn (Camacho et al. 2009). The plasmid pWP5-S18-ESBL-09_1 (NZ_AP022172.1) was reported in *E. coli* WP5-S18-ESBL-09 (Sekizuka et al. 2022). Lines in the plot correspond to local alignments where the color gradient depicts the proportion of identical nucleotides in the alignment. Colored rectangles in (A, B) correspond to annotated GenBank features such as coding sequences (CDSs) on the plus strand (blue) and minus strand (red). Annotated pseudogenes (grey) indicated below. **(B)** Results of the chaining algorithm for the same plasmid. Lines in the plot correspond to identified chains (or segments) where the color gradient depicts the mean sequence similarity of local alignments (in proportion of identical nucleotides). **(C)** Comparison of gene content in the large segmental duplication detected in the above plasmid. Connections between CDS in grey gradients indicate amino acid sequence identities of ≥30%. Likely paralogous genes are colored in the same shade. **(D)** Illustration of steps in the chaining algorithm. (I) The input to the core algorithm is the local alignment data from a replicon against itself (or between two distinct sequences). Colored rectangles show local alignments corresponding to unique putative duplications along the query and subject sequence (II) Pairwise gaps between local alignments are calculated for both query and subject sequences. (III) The scaled gap matrices for the query and subject sequences are merged into a weighted adjacency matrix, approximating collinearity among local alignments. (IV) Components with positive weights in the adjacency matrix are extracted, capturing local alignments linked by bridged gaps. (V) Local alignments connected in the network are chained together, resulting in newly defined coordinates for the detected segmental duplications.

A full detection of such segmental duplication is unlikely to be achieved by a local sequence similarity search tool. The BLAST algorithm extends the local alignment only as long as the scoring criteria of high sequence similarity are met. Segmental duplications that underwent sequence divergence likely result in multiple collinear, fragmented alignments. The gaps between such BLAST hits are characterized by low sequence similarity that terminates the BLAST alignment extension process. Thus, while local similarity tools can be used for preliminary detection of duplication events, they are not sufficient for inferring the most parsimonious segmental duplication events.

### Chaining local alignments in a nutshell

To infer diverged segmental duplication events, we developed a chaining algorithm that joins consecutive local alignment hits into a single continuous segment. Here we explain the fundamental principles of the chaining algorithm using an illustrative example of a plasmid genome. Initially, the plasmid is aligned to itself to generate local alignment hits, which serve as seeds for the chaining process. The hits are indexed and sorted based on their positions in the query sequence. Next, a pairwise distance is calculated between all alignment hits (Fig. 1D). This distance is used to detect gaps between local alignments that may stem from alignment extension abortion (and can be potentially bridged). Here we consider the gap size relative to the length of the flanking alignment hits. The distance metric *D*_*i*,*j*_ termed *scaled gap metric*, is defined as:

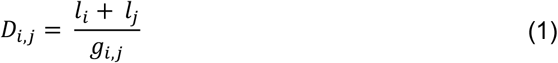

where:

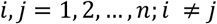

and relationship *i* and *j* must satisfy:

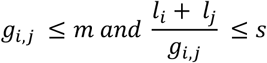

*D*_*i*,*j*_ describes the scaled gap distance metric between local alignments *i* and *j*. Thereby, *l*_*i*_ and *l*_*j*_ denote the length of the alignment hits with index *i* and *j*, while *g*_*i*,*j*_ is the absolute gap length separating them (in nucleotides). The thresholds *m* (maximum gap length) and *s* (maximum scaled gap) can be set for the chaining process to control the proportion of local alignments to gap length. These parameters help, for instance, to avoid chaining of consecutive, short local alignments that are widely spaced. This approach allows the algorithm to effectively identify and bridge fragmented alignments while avoiding overextension of the alignment in regions of low sequence similarity.

To compute the scaled gap metric for all local alignment pairs, we introduce scaled gap matrices for the query and subject sequences denoted as **Q** and **S** (Fig. 1D):

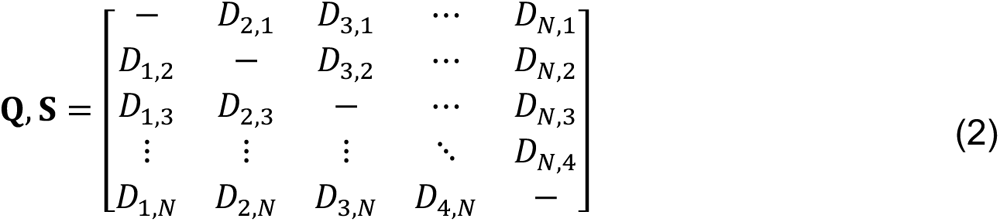

The query (**Q**) and subject (**S**) scaled gap matrices contain *D*_*i*,*j*_ metrics as N × N matrices, where N is the total number of local alignments. Each element *D*_*i*,*j*_ in the matrix corresponds to the scaled gap metric between alignment *i* and *j* (Fig. 1D). Entries along the diagonal (*D*_*i*,*j*_) correspond to self-comparisons hence are undefined. Entries where the scaled gap metric falls below the thresholds *m* and *s* are excluded (i.e., proportional control measurement of local alignment to gap length). To ensure proper comparability and chaining, **Q** and **S** store *D*_*i*,*j*_ metrics (i.e., scaled gaps) at the same matrix position, which also corresponds to the same alignment indices. This structured approach facilitates an efficient computation and ensures that the matrices accurately reflect the relationships between pairs of local alignments.

To combine the pairwise scaled gaps (i.e., proximity and alignment length) among local alignments along the query sequence with the corresponding alignments along the subject sequence, the algorithm merges the scaled gap matrices **Q** and **S** into a single weighted adjacency matrix **A**. Each element of **A** is defined as:

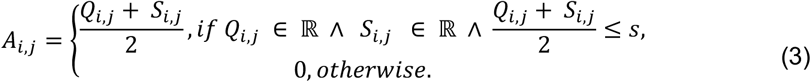

The resulting matrix **A** captures the pairwise scaled gap relationships between corresponding alignments in the query and subject sequences. Undefined relationships and scaled gaps that fall outside the specified threshold *s*, are modified to zero (Fig. 1D). To that end, connected local alignments in the adjacency matrix are considered network components that are chained into cohesive segments reflecting diverged, segmental duplications.

A common limitation in chaining algorithms is their applicability to circular replicons due to linear sequence representation in FASTA-formats. Circular topology, as in plasmid replicons, can disrupt the chaining process by failing to connect local alignments that span the contig start and end positions. To address this functionality, our method temporarily duplicates circular sequences during the chaining process, allowing seamless alignment connections. After chaining, segment coordinates that exceed the original sequence length are adjusted to their corresponding positions.

To enhance the tool usability and enable further customization, we structured SegMantX into five main modules, each addressing specific steps in the workflow (see Supplementary Figure 1). The output of the chaining process includes segment coordinates, indices of chained alignments, number of chained alignments, and metrics such as percentage of alignments to gap lengths, etc. The resulting data can be used to visualize identified segments interactively or to extract segment sequences for downstream analysis.

### Evaluation of SegMantX’s algorithm performance using simulated data

Under which conditions BLAST fails to infer continuous alignments (i.e., complete segmental duplications)? To answer that question, we introduced simulated sequence divergence in 21,452 randomly chosen plasmid genomes. Single-nucleotide mutations were introduced to a randomly chosen region of plasmid genomes, with variable mutated sequence length and mutation frequency (Supplementary Figure 2). Pairs of original and simulated plasmid sequences were aligned using BLASTn (Supplementary Figure 3). A total of 9,786 simulations (46%) produced fragmented alignments, which were most prevalent if the simulated diverged regions had ≤60% nucleotide sequence identity and were ≥75bp long (Supplementary Figure 4). Even 60% sequence identity in regions as long as 200 bp, or shorter regions (75 bp) with 40% sequence identity, frequently resulted in alignment fragmentation. The gap simulation results demonstrate that BLAST’s ability to produce continuous alignments is limited under specific divergence and length thresholds.

Using the same simulation procedure as described above, we evaluated the nucleotide local alignment module of MMSeqs2 (Steinegger and Söding 2017). In detail, MMSeqs2 yielded 8,473 (40%) discontinuous local alignments most of them being introduced with a simulated gap length ≥2500 bp (Supplementary Figure 5). Thus, sequence alignment settings required to yield complete alignments using MMSeqs2 are more dependent on the sequence length of the introduced divergence, compared to BLAST, which revealed clearer thresholds for alignment fragmentation. Notably, none of the parameter sets indicate a range where continuous alignments have been reliably retrieved using MMSeqs2. Thus, BLAST is superior for the purpose of chaining local alignments as the alignment settings yielding complete alignments for MMSeqs2 are less predictable.

To evaluate the chaining algorithm performance, we applied it to simulated plasmid sequences. Sequences in variable length were extracted from plasmid genomes and evolved using 10,000 random parameter sets that varied in total hit length (i.e., flanking regions), gap length (i.e., surrounded regions), hit sequence identity, and gap sequence identity (Fig. 2A). The chaining algorithm was subsequently applied to local alignments between the original and evolved (simulated) plasmid sequences, with a scaled gap metric threshold equal to 1. The results show that 412 (∼4%) of the simulated sequences were identified as continuous alignment by BLAST. Additional 1,736 (∼17%) simulated sequences were successfully aligned with the original sequence using the chaining algorithm. Comparing the alignment metrics of chained local alignments to continuous alignments detected by BLAST shows that the identity metrics (hit identity, gap identity, and global sequence identity) had a similar mode, yet the chaining algorithm was successful over a much wider range of sequence identity parameters (Fig. 2B-D). The median values for chained alignments were worse than those for BLAST-derived continuous alignments (see median lines in Fig. 2B-G). Sequences that were successfully chained had significantly higher global sequence similarity compared to those that remained discontinuous (P < 0.001, Wilcoxon test; global nucleotide sequence identity: median_chained_=67%, median_discontinuous_=44%).

**Figure 2.**
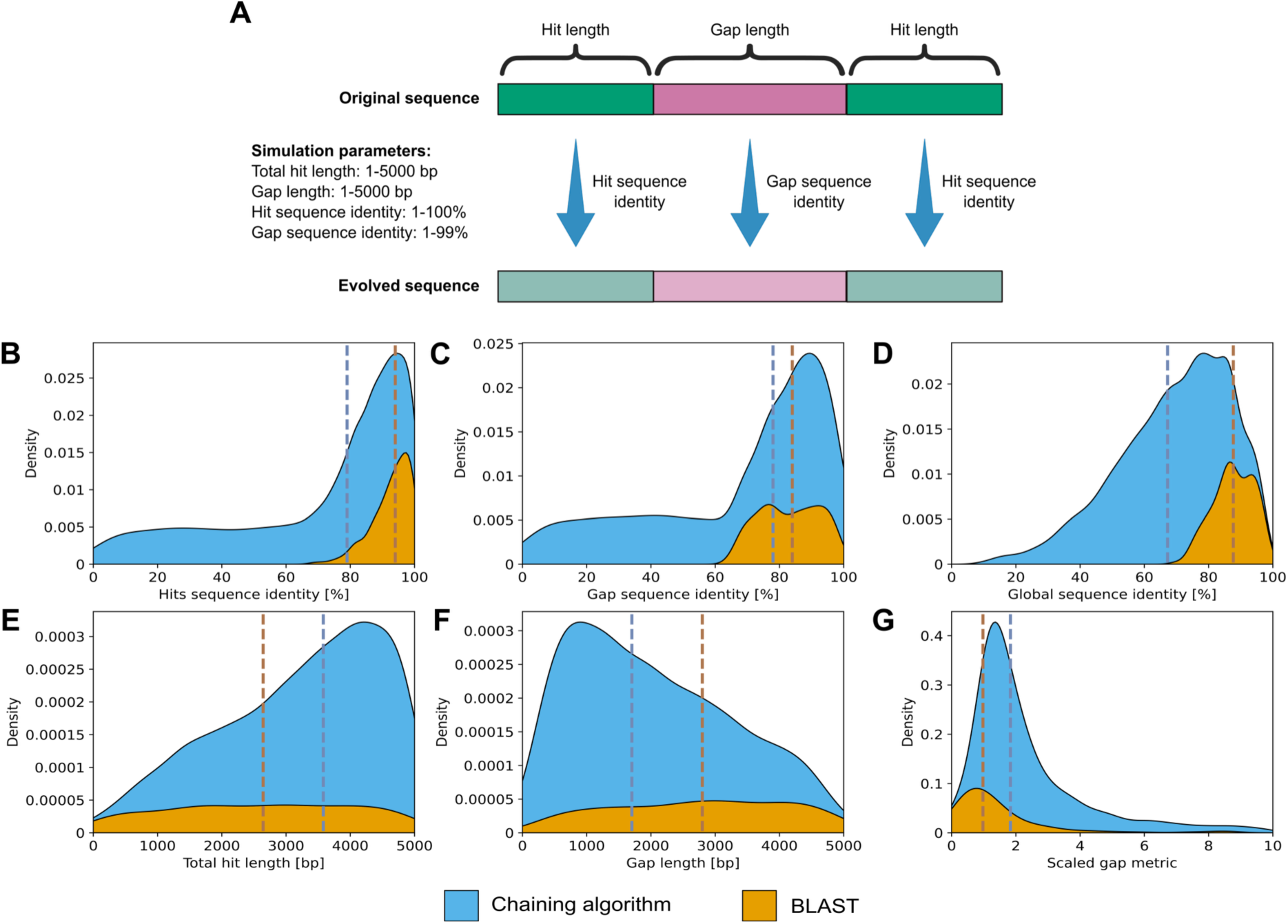
Performance of chaining local alignments compared to BLAST using simulated data. **(A)** Illustration of the simulation parameters. 10,000 sequences were extracted randomly from 1,741 plasmids that are characterized by no significant local alignments. The total length of the flanking hits, the gap length, hit sequence identity, and gap sequence identity have been randomly selected (within the stated parameter range). The length of the flanking regions and their sequence divergence is identical. The original and evolved sequences were aligned with BLASTn and the resulting local similarity hits were chained. **(B)** The density plots display the distribution of alignment metrics for complete alignments retrieved by BLAST (orange) and our chaining method (blue). The y-axis represents the normalized density, where the area under each curve has been scaled so that the total area across all curves is equal, allowing for a direct comparison of the relative densities across different distributions. Medians for the density distributions are represented as vertical dashed lines. For visualization purposes the x-axis for the scaled gap metric density plot was limited to ten (n=1,943 simulations, ∼90%).

Comparing the distribution of total hit length and gap length shows approximately uniform distributions for the continuous BLAST alignments (Fig. 2E & F). The chaining approach succeeded in aligning ca. 4-fold simulated sequences, and showed left-skewed total hit length distribution and right-skewed gap length distribution. This result reflects that BLAST performance is based on sequence identity, alignment score measurements, and alignment extensions regardless of the hit length and gap length (see also Supplementary Figure 6). In contrast, our chaining algorithm distinctly favors joining long, consecutive alignments separated by a smaller gap size (see also Supplementary Figure 7). In addition, BLAST yielded continuous segments below our applied scaled gap metric threshold (see *s* in Formula 1 and Fig. 2G) of one, but most of the continuous alignments retrieved by BLAST are commonly characterized by a high hit and gap sequence identity. Taken together, by applying our chaining algorithm to simulated data, we demonstrate its effectiveness for identifying robust consecutive local alignments and merging them into continuous segments, making it particularly well-suited for detecting segmental duplications.

### SegMantx reveals prevalent segmental duplications in enterobacterial plasmids

Applying the chaining algorithm to all enterobacterial plasmids in our dataset recovered 52,553 segments in 4,400 plasmids (64%). Of these, 27,038 (51%) were the result of chaining, and the remaining 25,515 (49%) segments could be detected directly by BLAST. Plasmids lacking detectable duplications are significantly smaller in size (P < 0.001, Wilcoxon test; median size with duplications = 97,839.5 bp; without duplications = 5,167.5 bp) and encode substantially fewer coding sequences (CDSs) (P < 0.001, Wilcoxon test). The segments recovered by chaining were significantly larger than those obtained from BLAST alone (P < 0.001, Wilcoxon test; median chaining segment = 823 bp, median BLAST segment = 677 bp) where majority of chained segments include CDSs (79%).

To further explore the frequency of duplication events in plasmid genomes, we define four categories of duplication events (similarly to previous described classifications, Graur et al. 2016; Panchy et al. 2016; Almeida-Silva and Van De Peer 2025) (see Fig. 3A and methods). Here we demonstrate the four categories using example duplications inferred with SegMantX. Duplicated genes can be organized in tandem or dispersed from one another. A tandem duplication of ∼6.6 kb was inferred in a *S. enterica* conjugative plasmid (Fig. 3B). The two segments in the duplicated region include two conserved and two homologous gene functions (see Fig. 3F and Supplementary Table 2). A dispersed duplication was inferred in a *K. pneumoniae* plasmid; the duplicates located ∼18 kb apart, include five conserved genes (Fig. 3C & F, Supplementary Table 3). Gene duplications associated with mobile genetic elements (MGEs) were identified as such if they included MGE-associated gene families in the *KES* dataset (see Methods). Example is a *K. pneumoniae* plasmid where we inferred a duplication spanning ∼3.6 kb that includes two conserved genes annotated as Tn3-like family transposases and recombinase (Fig. 3D & F, Supplementary Table 4). The fourth category is a plasmid homomultimer, that is, plasmids whole genome duplication events – that we term here whole plasmid duplications (WPDs). Example is an *E. coli* plasmid pU90-2. The duplication spans the entire 18,701 bp plasmid genome, with two 9,050 bp duplications, each encoding eleven CDSs typical of ColE1 plasmids (e.g., Rop, colicine, and mobilization-related genes, see Supplementary Table 5). Notably, one IS66 family-like insertion sequence within the duplication has 5% amino acid sequence divergence, while the other genes are 100% conserved, which may suggest a recent WPD event.

**Figure 3.**
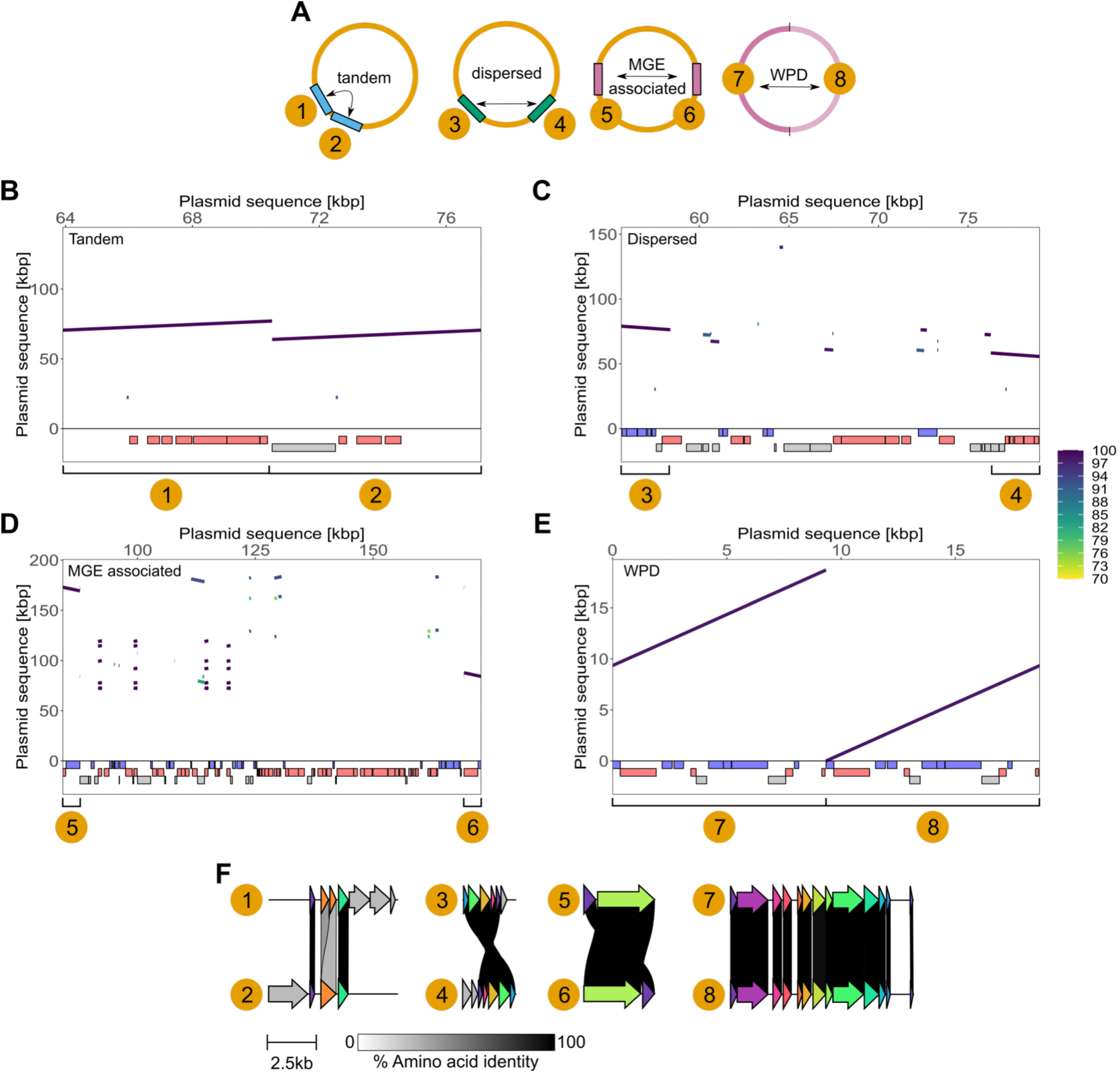
Classification of segmental duplications into tandem, dispersed, or MGE-related gene duplications and plasmid multimers. **(A)** Illustration of four types of duplication events identified in our dataset. **(B-E)** Zoomed-in views of the inferred duplications, highlighting specific duplication events in plasmids, are depicted. **(B)** pUnnamed2 (NZ_CP022035.1) in pathogenic *S. enterica* subsp. enterica serovar Onderstepoort SA20060086 (Robertson et al. 2018). **(C)** pRHB30-C05_3 (NZ_CP057315.1) in *K. pneumoniae* RHB30-C05 isolated from livestock (AbuOun et al. 2021). **(D)** Antibiotic resistance plasmid pKpN06-CTX (NZ_CP012993.2) in *K. pneumoniae* KpN06 isolated from human (Lynch et al. 2016). **(E)** pU90-2 (NZ_CP068037.1) in *E. coli* U90 hosted by swine with gastroenteritis. Lines in the plot correspond to local alignments or segments where the color gradient depicts the mean sequence similarity of local alignments (in proportion of identical nucleotides). Colored rectangles in (B, C, D, E) correspond to annotated GenBank features: coding sequences (CDSs) on the plus strand (blue) and minus strand (red). Annotated pseudogenes (grey) indicated below. Brackets below each panel highlight the inferred segmental duplications. **(F)** Comparison of gene content in the inferred segmental duplications (see also numbers in B-E). Connections between CDS in grey gradients correspond to amino acid sequence identities of ≥30%. The color scheme used for CDS corresponds to homologous amino acid sequences.

### Most segmental duplications in plasmids are associated with mobile genetic elements

Which type of duplications are common in plasmid evolution? To that end, we classified the 52,524 inferred duplications into the four categories, tandem, dispersed, MGE-associated and WPD (see Methods and examples in Fig. 3 B-E). The segments were further sorted according to their annotations and length (in genes) into non-coding duplications, single gene duplication, segmental gene duplication (i.e., ≥2 genes), and WPDs. Large duplications such as segmental gene duplications and plasmid multimers had a low frequency compared to non-coding and single gene duplications (Fig. 4A). Most of the 16,789 (84%) single gene duplications correspond to prominent MGE-related gene functions, while 5,709 (84%) of the segmental duplications include insertion sequences (ISs) and transposases. Consequently, we conclude that MGE-related functions account for the largest fraction of duplicated genes in plasmid genomes.

**Figure 4.**
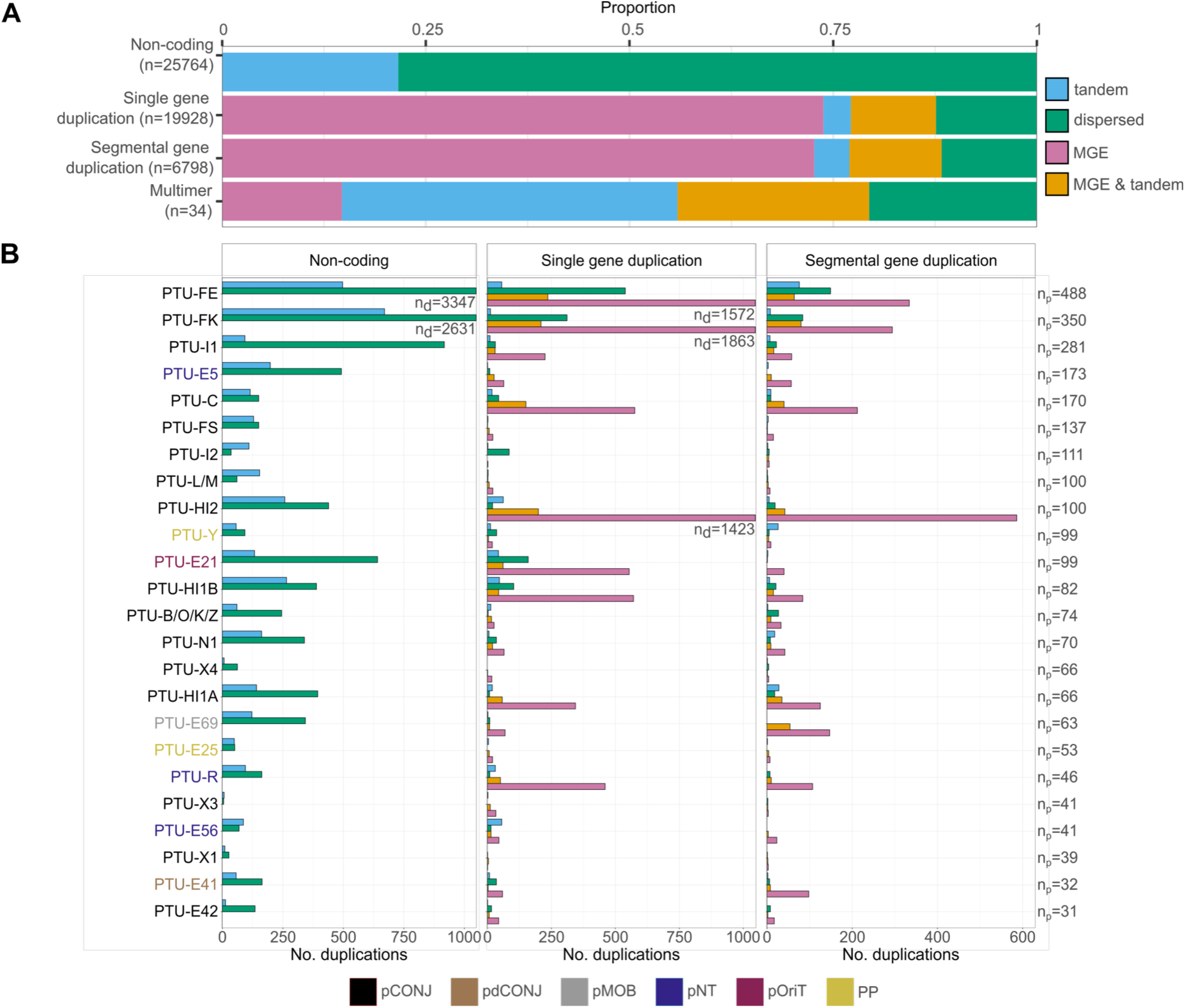
MGE-associated duplications are prevalent in all PTUs. **(A)** Proportions of duplication types across plasmid genomes. **(B)** Duplication classification and frequencies per PTU. The PTU labels coloring depict the main plasmid mobility types classified as conjugative (pCONJ), decayed conjugative (pdCONJ), mobilizable (pMOB), non-transmissible (pNT), phage-plasmids (PP), and *oriT*-bearing plasmids (pOriT). Note, that the graph includes a proportion of the *KES* plasmids (n=2,688, 61%) that are characterized by duplications and could be assigned to their mobility type mobility classifications as per Ares-Arroyo et al. 2024. PTUs that are included in the graph correspond to plasmid groups that are characterized by larger plasmid size in the *KES* dataset (Wang and Dagan 2024). The labels n_d_ indicate the number of duplications exceeding the x-axis truncation, and n_p_ represents the number of plasmids within each PTU.

To further examine whether duplication frequency is similar among plasmid lineages, we compared the prevalence of the four duplication classes among related plasmid groups in *KES* inferred from plasmid taxonomic units (PTUs) (Redondo-Salvo et al. 2020; Redondo-Salvo et al. 2021). Duplications of non-coding (and dispersed) regions is the most frequent duplication class for the majority of PTUs (n=18, 75%) (Fig. 4B). Single gene duplications associated with MGEs in all PTUs are prevalent (and redundant). In 21 (88%) PTUs the most frequent category of single gene duplications corresponds to MGEs. Segmental gene duplications are rather rare among all PTUs, but if present, they are commonly associated with MGEs as well. PTU-Y, which comprises phage-plasmids, is an exception where tandem segmental duplications are more frequent than single-gene duplications. These duplications often encode phage-related functions, including tail fiber, phage tail, holin, and recombinase family proteins, potentially associated with gene amplification in such plasmids. Furthermore, only two WPDs were uncovered among PTUs corresponding to plasmid homomultimers in conjugative PTU-X1 (NZ_LS999563.1) and PTU-X3 (NZ_CP025215.1) which are plasmids groups with medium plasmid size averages of 30kb and 44kb (see also Peñil-Celis et al. 2024). Our findings highlight an important role of MGEs in plasmid gene duplication.

### Combinations of co-duplicated functions match MGEs and their antibiotics resistance genes cargo

Our approach for the inference of segmental duplications has the advantage of recovering duplication events comprising multiple genes. To that end, we asked which gene functions are commonly duplicated and to what extent they are retained. Therefore, we classified all gene families of the *KES* dataset into functional categories. The majority of duplicated gene families in *KES* plasmids correspond to “Transposable elements and IS”, “Hypothetical”, and “AMR” followed by the categories “Replication, recombination, and repair,” and “Transcription”, which are mostly MGE-related as well, e.g., integrases, recombinases, and relaxases (Fig. 5A).

**Figure 5.**
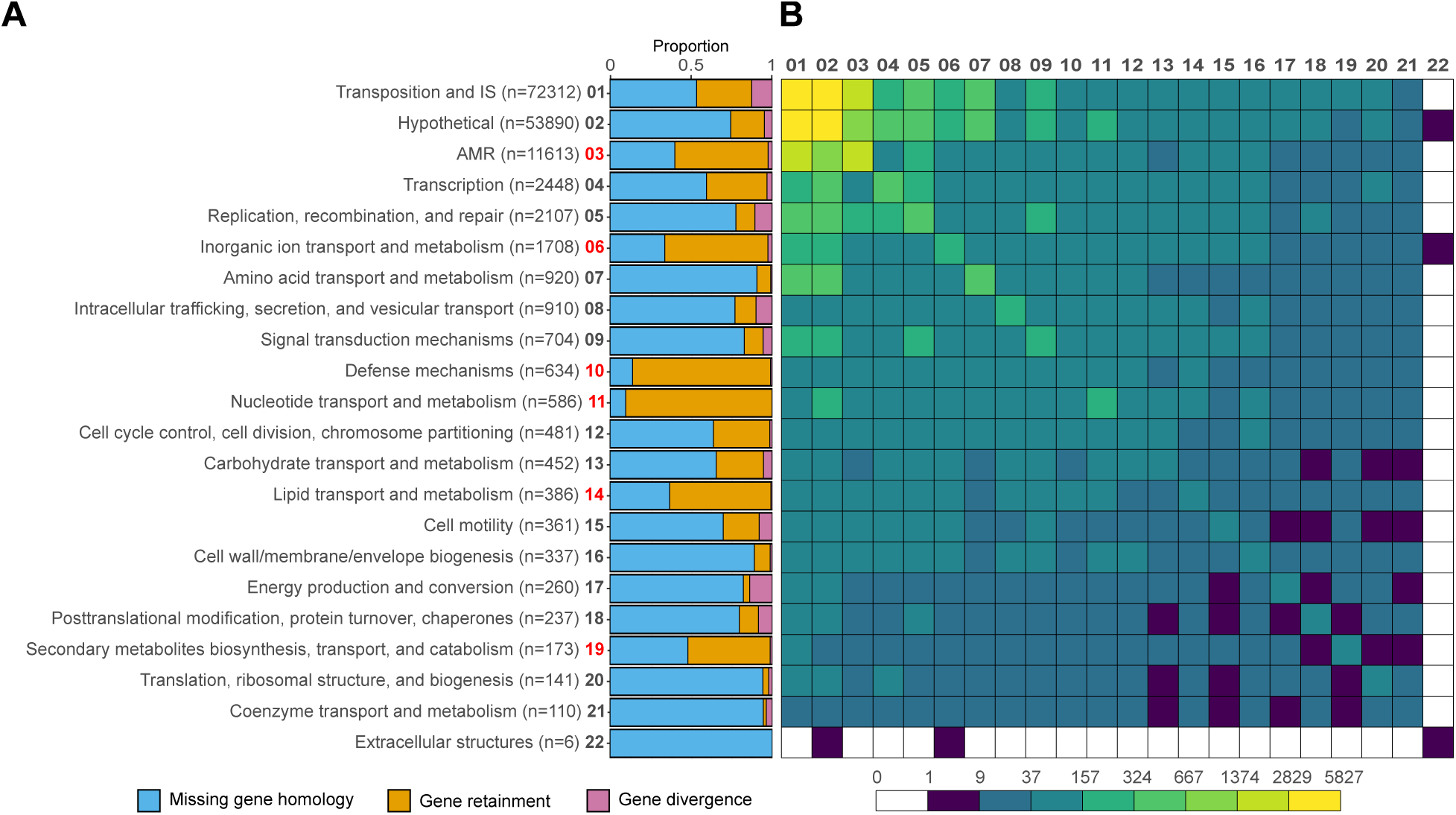
Co-transferred gene functions in plasmid duplications are functionally related. **(A)** Proportions of gene retainment categories for CDS based on functional annotation. Mobile genetic elements, hypothetical proteins, and antibiotic resistance genes (AMRs) were classified according to gene annotations, supplemented by a thorough gene family classification of AMRs and MGE-related functions (Wang and Dagan 2024). The x-axis labels include a sequential index integer where each functional category is listed in panel (B). Red x-axis labels correspond to functional annotation categories where gene retention shows a higher proportion than missing gene homologies. **(B)** Heatmap displaying the frequency of paired gene function combinations observed in segmental plasmid gene duplications. The heatmap axes correspond to the functional annotation categories in (A). The logarithmic color gradient in the heatmap shows the frequency of paired gene functions found in segmental duplications (see legend for scale).

To further evaluate which functional categories are retained, we determined orthologous genes between duplicates to calculate the level of their sequence conservation (as % of identical amino acids). Most of the duplicated functional categories show absence of homology between duplicated regions (n= 91,396, 61%). However, some exceptions reveal a major proportion of retained genes in functional categories (see red labels in Fig. 5A). Note, that our method does not estimate the time since duplication event, which may further indicate the fate for gene duplications in the long-term, that is, highly conserved duplicated genes may correspond to recent duplication events. Gene divergence resulting from gene duplications (i.e., amino acid sequence identity < 100%, n = 12,618, 8%) further indicates a minor level of gene divergence with a median sequence identity of 98.5%. Overall, the absence of CDS homology and the retention of genes between duplicated regions are prominent in plasmid genomes, suggesting gene loss or gain as possible fates for gene duplicates, with retained functions likely resulting from recent duplication events.

To further evaluate whether segmental gene duplications comprise functionally related genes (i.e., operon-like organization), we analyzed the co-occurrence of functional categories in the inferred segments. The distribution of co-duplicated CDSs depicts an apparent high frequency on the diagonal indicating that genes classified into the same category tend to be duplicated together (Fig. 4B). This trend likely corresponds to the duplication of genes that are clustered together as functional unit (as in operons). Notably, the highest frequencies of co-duplicated gene functions were observed among combinations of “Transposition and IS”, “Hypothetical”, and “AMR” functional units, which are common combinations in MGEs, such as transposons and integrons. Transcription-, replication-, repair, and recombination-related genes are further frequent co-duplicated gene functions and are often duplicated alongside genes from other functional categories related to transposition, unknown functions, and metabolism (Fig. 5B). These findings highlight the potential role of segmental gene duplications in maintaining or amplifying functional clusters, which may facilitate adaptive evolution and the spread of advantageous traits within mobile genetic elements and across genomes.

### Beyond DNA duplication inference: leveraging local alignment chaining to identify DNA transfer

To extend the applicability of our tool, we have adapted our chaining approach to enable local alignment chaining between two distinct sequences. First, we tested our modified method by chaining BLASTn results between plasmid pTEF1 and the chromosome in *Enterococcus faecalis* V583, following recent reports of gene sharing in this genome (Sanchez-Herrero et al. 2020; Kadibalban et al. 2024). Briefly, SegMantX identified fewer but larger segments (808-21,680 bp) compared to a hit-agglomeration strategy (Kadibalban et al. 2024), demonstrating that the chaining approach yields less fragmented segments (Fig. 6A). The largest segment in pTEF1 corresponds to 21 CDSs mainly including transfer-, replication-related, and hypothetical proteins. Eight plasmid segment CDSs are identical, and 12 show homologies (≥70% amino acid sequence similarity) to CDSs within the chromosomal segment, maintaining a syntenic gene order (Fig. 6B, Supplementary Table 6). Applying SegMantX to chain local alignments between pTEF1 and the chromosome in *E. faecalis* V583, yielded a 21,680 bp cohesive segment sharing with syntenic gene order. This pattern of shared sequence similarity likely corresponds to a past DNA transfer event that would have not been fully reconstructed using other approaches (e.g., BLASTp or hit-agglomeration strategy, Sanchez-Herrero et al. 2020; Kadibalban et al. 2024).

**Figure 6.**
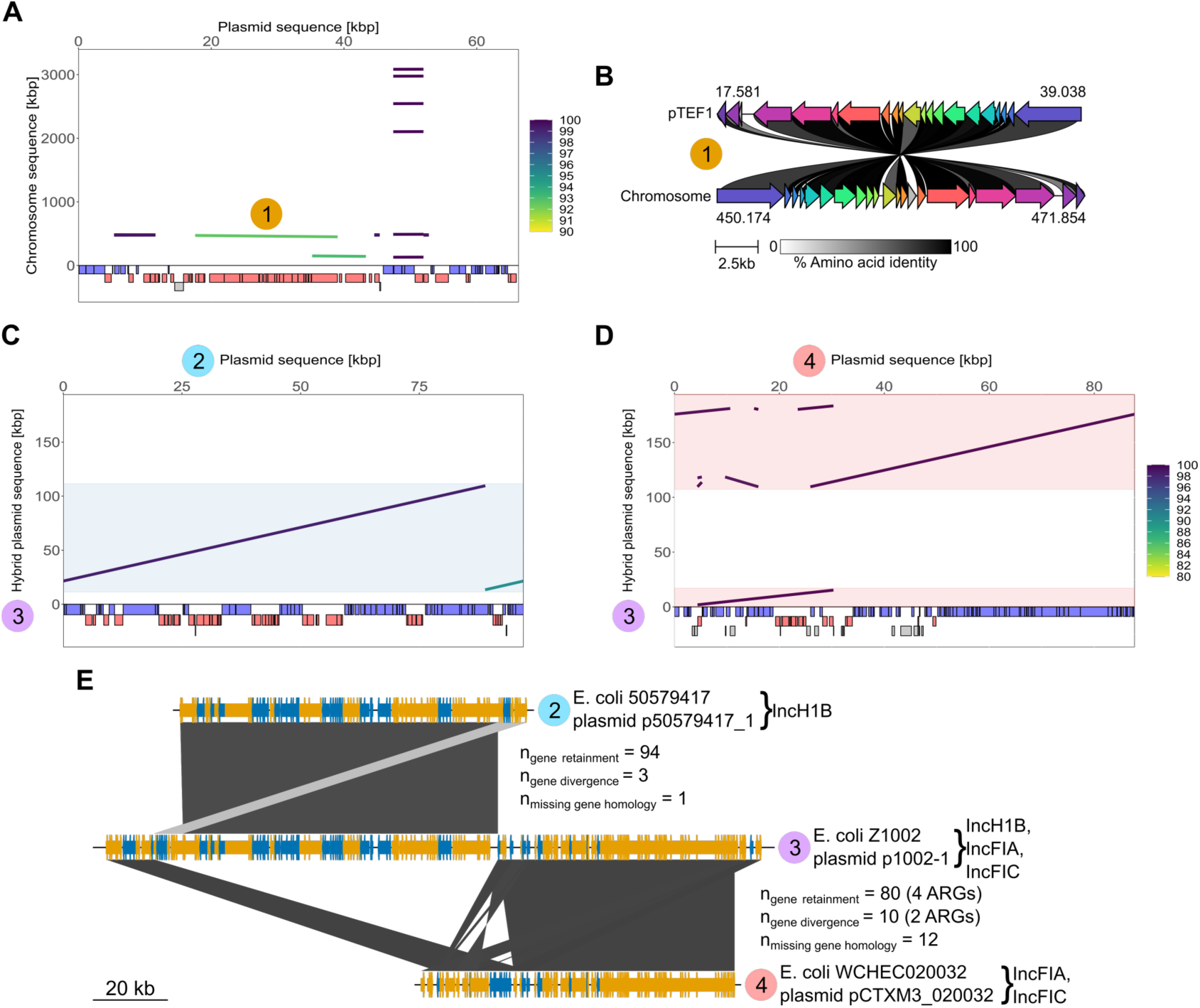
Detection of DNA transfer between plasmids and chromosomes and plasmid fusions. **(A)** Results of the chaining algorithm for plasmid pTEF1 against the chromosome in *Enterococcus faecalis* V583 (GCF_000007785.1). Lines in the plot correspond to identified chains (or segments) where the color gradient depicts the mean sequence similarity of local alignments (in proportion of identical nucleotides). Colored rectangles in (A, C, D) correspond to annotated GenBank features: coding sequences (CDSs) on the plus strand (blue) and minus strand (red). Annotated pseudogenes (grey) indicated below. The circle with a serial number indicates the segment whose gene content was compared in (B). **(B)** Comparison of gene content in the inferred shared DNA between plasmid pTEF1 and *E. faecalis* V583 chromosome (see also labeled segment in A). Connections between CDS in grey gradients correspond to amino acid sequence identities of ≥30%. The color scheme used for CDS corresponds to homologous amino acid sequences. **(C, D)** Results of the chaining algorithm for plasmids p50579417_1 (C) in *E. coli* 50579417 (GCF_003813145.1) and pCTXM3_020032 (D) in *E. coli* WCHEC020032 (GCF_002164865.2) against plasmid p1002-1 in *E. coli* Z1002 (GCF_002142675.1). Lines in the plot correspond to identified chains (or segments) where the color gradient depicts the mean sequence similarity of local alignments (in proportion of identical nucleotides). Colored rectangles (blue, red) in the plots highlight plasmid genome regions of p50579417_1 and pCTXM3_020032 (x-axes) that are aligned to the hybrid plasmid p1002-1 (y-axis). The circles with a serial number indicate the plasmid sequences whose chaining results were compared in (E) using genoPlotR (Guy et al. 2010). **(E)** Results of our chaining algorithm (i.e., segments) for plasmids compared in (C, D) using genoPlotR (Guy et al. 2010). Arrows represent CDSs on the plus (orange) and minus (purple) strand. Amino acid sequence alignments between plasmids have been categorized into gene retainment (AA_identity_=100%), gene divergence (AA_identity_<100% and ≥30%), and missing gene homology (AA_identity_<30%) based on BLASTp results using 1e-9 as E-value cutoff.

To further demonstrate the range of SegMantX’s use cases, we present its application for the detection of hybrid plasmids (i.e., multi-replicon plasmids). Analysis of the conjugative phage-plasmid p1002-1 revealed that this plasmid sequence may correspond to a hybrid replicon composed of conjugative plasmid pCTXM3_020032 and phage-plasmid p50579417_1, all of them residing in closely-related *E. coli* isolates (Fig. 6C, D). Comparing CDSs of the conjugative and phage-plasmid to the hybrid plasmid, shows a similar gene composition. Furthermore, the conjugative plasmid encodes six antibiotic resistance genes with homologous CDSs in the hybrid plasmid (Fig. 6E). The conserved syntenic gene and segment order detected by SegMantX support the notion that p1002-1 evolved via an ancient plasmid fusion event (Fig. 6E). The fusion event thus led to the emergence of a conjugative phage-plasmid.

## Discussion

SegMantX provides a framework for chaining sequence alignments, particularly towards fragmented duplication events following sequence divergence. Introducing a threshold metric that balances local sequence alignment proportional to the gap length furthermore provides a quantitative method for identifying consecutive similarity hits that likely stem from genuine homologous regions. Evaluating the performance of our approach demonstrated that traditional local alignment methods could not detect these patterns due to sequence divergence post-duplication event (e.g., BLAST, MMSeqs2, Camacho et al. 2009; Steinegger and Söding 2017). We adapt the concept of chaining local sequence alignments (e.g., DAGChainer and YOC, Haas et al. 2004; Uricaru et al. 2015) to demonstrate that using sequence similarity searches from conservative tools like BLASTn as seeds can yield continuous alignment parameters with well-defined boundaries to discontinuous alignments (Camacho et al. 2009). In contrast, when using MMSeqs2 to generate local alignments as seeds for the chaining process, the resulting boundaries distinguishing continuous from discontinuous alignments were less reliably defined (Steinegger and Söding 2017). SegMantX can process seed data for chaining using other approaches, as our method relies on alignment positions in sequences. The implementation of SegMantX both as command-line tool for batch processes and user-friendly graphical interface enable an interactive use of the tool for validation purpose, as well as its integration in large-scale workflows. Previous gene duplication detection tools may have limitations in usability, or fail to account for transposon-derived duplications, that restrict the scope of their inference (e.g., MCScanX, DupGen_finder, Wang et al. 2012; Qiao et al. 2019). We demonstrate that SegMantX performs well in chainining local alignments between two distinct sequences, for instance, towards inference of gene transfer. Demonstrating plasmid hybridization and gene transfer between a chromosome and plasmid highlights SegMantX as an alternative approach to existing strategies for sequence homology inference.

Applying SegMantX to enterobacterial plasmids reveals a low frequency of large-scale segmental duplications and whole-plasmid duplications (WPDs). The rare whole plasmid duplication events reported here (n=34; 0.05%), were often not affiliated with a plasmid taxonomic unit (PTU). The absence of a PTU classification may stem from limited sampling density or the inherent technological limitation of genome sequencing for detection of WPDs, which relies on the read length and quality of the sequencing (Juraschek et al. 2021). Alternatively, our results suggest that plasmid homomultimers are generally transient as they are expected to be outcompeted by plasmid monomers (e.g., under non-selective conditions, Wein et al. 2020), with only a subset persisting long enough to expand into a detectable plasmid lineages that can be taxonomically classified. Genes in the duplicated region may become over time non-functionalized or lost, as observed in chromosomal duplications from *Salmonella* evolution experiments (Cao et al. 2022). The large-scale segmental duplications in *KES* plasmids appear to be recent, transient events that primarily result in an immediate gene dosage effect but are rarely retained. In the absence of strong positive fitness effect of the segmental duplication, rare plasmid alleles are expected to be rapidly lost due to segregational drift (Garoña et al. 2023). Large structural duplications in plasmids are therefore more likely to be fixed if (some) duplicated genes initially provide the host a fitness advantage.

Here we classify the inferred duplications using a canoncial approach into tandem, dispersed, MGE-mediated, whole-plasmid duplications, similarly to other available tools (e.g., MCScanX-transposed or doubletrouble, Wang et al. 2013; Almeida-Silva and Van De Peer 2025). The detection of gene duplication with SegMantX does not enable a distinction between paralogs and xenologous, which could be performed in a downstream analysis (e.g., using phylogenetics and gene presence/absence patterns, Tria and Martin 2021). Indeed, the role of mobile genetic elements (MGEs) in gene duplication blurs the line between paralogs and xenologs. Genes associated with MGEs are typically considered xenologs, yet, MGEs act as intermediaries in both gene duplication (i.e., within the same replicon) as well as gene transfer, e.g., between species (but also between distinct replicons). For example, a study of *Staphylococcus* genome evolution showed that MGEs not only facilitate gene transfer but also contribute to gene amplification (Chan et al. 2011). Duplications driven by MGEs may thus be considered as ‘synologs’, a term that merges paralogs and xenologs into a single class of homologous genes within a genome (Lerat et al. 2005).

Our study reveals MGE-associated duplications as the most frequent category in plasmid evolution, in agreement with previous studies showing that gene duplications in prokaryotes are often associated with transposons, retrotransposons, and other mobile genetic elements (e.g., Tria and Martin 2021). This result is also consistent with our recent study showing frequent non-functionalization events of mobile genetic elements that proliferate in plasmids (Hanke et al. 2024). Hence we infer that segmental duplications in plasmids are likely associated with high gene turnover, in aggreement with previous findings (Vos et al. 2015). Our results highlight SegMantX as a useful framework to detect segmental duplications without prior assumptions on the fate of duplicated genes.

## Materials & Methods

### Genomes data and gene families

The *KES* dataset, adopted from (Wang and Dagan 2024), comprises 1,114 chromosomes and 3,098 plasmids from *Escherichia*, 755 chromosomes and 2,693 plasmids from *Klebsiella*, and 572 chromosomes and 993 plasmids from *Salmonella*. Protein-coding genes in the *KES* genomes are clustered into 32,623 gene families, as described previously (Wang and Dagan 2024).

### Determining putative duplicated regions using local sequence similarity searches

Putative DNA duplications in *KES* plasmids (n=6,784) were initially inferred with BLAST+ (v.2.14.0+, Camacho et al. 2009). Therefore, we created nucleotide BLAST databases of each plasmid sequence using makeblastdb with parameter -dbtype nucl. Subsequentially, BLASTn was used to perform a local sequence similarity search of each plasmid sequence against itself using blastn with parameters -num_threads 1 -outfmt 7 -evalue 1e-9 -perc_identity 60 -dust no - soft_masking F. Local sequence similarities identified by BLASTn searches were analyzed to detect potential DNA duplications.

### Chaining local alignments using SegMantX towards DNA duplication detection

To identify DNA duplications, we first performed local similarity searches of *KES* plasmids against themselves using SegMantX’s module generate_alignments with parameters -q [plasmid_fasta_file] -Q -SA -i 60 -e 1e-9 -a [alignment hits output file]. The resulting self-sequence alignment output data of individual plasmids was the input for the chaining process using SegMantX’s module chain_self_alignments with parameters -i [alignment hits output file] -G 5000 -SG 1 -f [plasmid fasta file] -ml 200 -Q -o [chaining output file]. The resulting nucleotide chains (or segments) have been extracted for all plasmids using SegMantX’s module fetch_nucleotide_chains with parameters -i [chaining output file] -fq [plasmid fasta file] -o [chains output fasta file]. Dotplots of the resulting segments were generated using SegMantX module visualize_chains with parameters -i [chaining output file] -S kbp -fq [plasmid fasta file] -gf [plasmid GenBank file] -QIS -o [plot output file]). Our chaining tool is based on python and was developed for command line application or using a graphical user-interface based on streamlit. For further documentation, installation, and examples visit: https://github.com/DMH-biodatasci/SegMantX.

The comparison and visualization of gene content in duplicated regions was done using clinker (Gilchrist and Chooi 2021) (v0.0.29, default settings) with GenBank files trimmed to include only the detected duplications.

### Simulating plasmid genome evolution

To evaluate the conditions under which BLASTn or MMSeqs2 fail to detect continuous alignments, we aimed to introduce single-nucleotide mutations into randomly chosen regions of plasmid genomes, varying in mutated sequence length and mutation frequency to its original plasmid genome sequence (Supplementary Figure 2). A random sequence divergence ‘gap’ was simulated per randomly selected plasmid genome of the *KES* dataset resulting in 21,452 mutated plasmid genomes. Thereby, plasmid sequence mutation simulations were constructed by randomly sampling a plasmid sequence from 1,741 *KES* plasmids that did not reveal any significant local alignments in the BLASTn similarity search. Original and simulated (i.e., evolved) plasmid sequences were then aligned using BLASTn (v.2.14.0+, Camacho et al. 2009) with parameters -num_threads 1 -outfmt 7 -evalue 1e-9 -perc_identity 60 -dust no -soft_masking F or MMSeqs2 easy-search module (v.13.45111, Steinegger and Söding 2017) with parameters -e 1e-9 –min-seq-id 0.6 --search-type 3. Afterwards, the alignment continuity between original plasmid sequences and evolved plasmid sequences has been assessed for BLASTn and MMSeqs2 separately.

To evaluate the chaining algorithm performance, we randomly sampled 10,000 subsequences from 1,741 *KES* plasmids of the dataset that did not reveal any significant local alignments in the BLASTn similarity search. We further defined random parameter sets that varied in total hit length, gap length, hit sequence identity and gap sequence identity to generate sequences of simulated sequence divergence compared to their original plasmid subsequences (see also Fig. 2A). Afterwards, using the 10,000 diverged subsequences and their original sequence counterparts, we assessed the chaining performance measuring the following scenarios: 1) continuous alignment using BLASTn stand-alone (v.2.14.0+, blastn with parameters-num_threads 1 -outfmt 7 -evalue 1e-9 -perc_identity 60 -dust no -soft_masking F), 2) continuous alignment recovered by our chaining method (SegMantX’s module chain_alignments with parameters -i [alignment output file] -G 5000 -SG 1 -fq [simulated plasmid fasta file] -fs [fasta file subject] -o [chaining output file]), and 3) no significant alignment or discontinuous alignment.

### Classification of plasmid duplications

The inferred duplications were classified into non-coding duplications (no CDS), single-gene duplications (1 CDS), segmental gene duplications (>1 CDS) according to the number of CDS in the duplication region. Whole-plasmid duplications (WPDs) were identified as duplicates that span the whole plasmid genome. Putative WPD events were manually inspected and further validated based on typical plasmid backbone gene content (e.g., plasmid replication-, transfer-, partitioning-, stabilization- and maintenance-related genes). Segmental duplications found with ≤2 CDSs between them were further classified as tandem duplications. MGE-associated duplicated regions were classified as such if they contained MGE-associated gene functions (see Methods section ‘Classification of duplicated genes into functional categories’). All other duplicates were classified as dispersed duplicates.

### Classification of duplicated genes into functional categories

To classify *KES* gene families into clusters of orthologous groups (COGs) we applied EggNog-mapper to most frequent amino acid sequence variant per gene family (Cantalapiedra et al. 2021) (v. 2.1.12, with module download_eggnog_data and emapper and parameters: -i –output_dir – mode diamond). Furthermore, duplicated gene families have been assigned to custom functional annotations ‘AMR’ and ‘Transposition and IS’ according to a thorough gene family classification of *KES* focusing on antibiotic resistance genes (ARGs) and MGEs (Wang and Dagan 2024). In addition, we assigned remaining gene families into the functional annotations ‘Transposition and IS’, ‘AMR’, and ‘Hypothetical’ based on NCBI’s Prokaryotic Genome Annotation Pipeline (Li et al. 2021).

### Inferring the fate of duplicated genes

To infer the fate of genes in duplicated segments, CDSs along the two duplicated were first paired by searching for reciprocal best hits (RBHs) between the duplicated segments using MMseqs2 easy-rbh module (v.13.45111, Steinegger and Söding 2017) with a threshold of *E*-value ≤ 1 × 10^−10^). The identified RBHs were further aligned with the Needleman-Wunsch algorithm using parasail-python (v. 1.2.4, Daily 2016). RBHs and genes within duplicated regions were classified into three categories: gene retention (100% amino acid sequence similarity), gene divergence (<100% and ≥30% amino acid sequence similarity), and gene loss (if no RBH was found).

### Plasmid typing

Plasmids were assigned to plasmid taxonomic units (PTUs) using COPLA supplying taxonomic information of their hosts (Redondo-Salvo et al. 2021). PTUs correspond to a classification scheme of closely related plasmids as inferred from average nucleotide identity (ANI) networks of whole plasmid sequences (Redondo-Salvo et al. 2021). Plasmid mobility types were inferred using a previous classification scheme into conjugative (pCONJ), decayed conjugative (pdCONJ), mobilizable (pMOB), non-transmissible (pNT), phage-plasmids (PP), and *oriT*-bearing plasmids (pOriT) (Ares-Arroyo et al. 2024). Incompatibility (Inc) groups of *KES* plasmids have been identified, as previously described (Wang and Dagan 2024).

### Chaining local alignments using SegMantX towards DNA transfer and plasmid fusion detection

We used plasmid pTEF1 and the chromosome in *Enterococcus faecalis* V583 (GCF_000007785.1) and *KES* plasmids (p50579417_1 in *E. coli* 50579417 (GCF_003813145.1), and pCTXM3_020032 in *E. coli* WCHEC020032 (GCF_002164865.2) against plasmid p1002-1 in *E. coli* Z1002 (GCF_002142675.1)) to demonstrate SegMantX’s utilities towards identifying DNA transfer and plasmid fusion events. We first performed local similarity searches using SegMantX’s module generate_alignments with parameters -q [fasta_file_query] -s [fasta_file_subject] -Q -S -i 60 -e 1e-9 -a [alignment hits output file]. The resulting sequence alignment output data was the input for the chaining process using SegMantX’s module chain_alignments with parameters -i [alignment hits output file] -G 5000 -SG 1 -fq [fasta_file_query] -fs [fasta_file_subject] -ml 100 -Q -S -o [chaining output file]. Dotplots of the resulting, flattened segments (for 2D presentation) were generated using SegMantX’s module visualize_chains with parameters -i [chaining output file] -S kbp -fq [fasta file query] -fs [fasta file subject] -gf [GenBank file] -o [plot output file]).

The comparison and visualization of gene content in the DNA transfer of pTEF1 and the chromosome of *Enterococcus faecalis* V583 (GCF_000007785.1) was done using clinker (Gilchrist and Chooi 2021) (v0.0.29, default settings) with GenBank file trimmed to include only the largest DNA transfer segment.

We further visualized the results of our chaining algorithm for plasmids p50579417_1 and pCTXM3_020032 against plasmid p1002-1 using genoPlotR (Guy et al. 2010). Afterwards, similarities of amino acid sequences encoded by the plasmid sequences have been assessed using blastp with parameters -num_threads 1 -outfmt 7 -evalue 1e-9 (v.2.14.0+, Camacho et al. 2009). Amino acid sequence alignments between plasmids have been categorized into gene retainment (AA_identity_=100%), gene divergence (AA_identity_<100% and ≥30%), and missing gene homology (AA_identity_<30%).

### Statistical tests

All statistical tests were performed using R (v.4.0.3). P-values below the function threshold (eps = 2.2x10^-16^) are reported as P < 0.001.

## Acknowledgements

We thank Barbara Cania, Lisa Hartmann, Johannes Effe, and Mario Santer for critical comments on the manuscript and SegMantX. This research was supported in part through high-performance computing resources available at the Kiel University Computing Centre.

## Author contributions

D.M.H. and T.D. conceived the study. D.M.H. designed and developed SegMantX. D.M.H. designed and performed the data analysis and visualizations. D.M.H. and T.D. interpreted the results and wrote the manuscript.

## Data availability

The data are provided in electronic supplementary material and SegMantX GitHub page: https://github.com/DMH-biodatasci/SegMantX.

