## Supplementary Material for "SegMantX: a novel tool for detecting DNA duplications uncovers prevalent duplications in plasmids"

#### Supplementary Tables

**Supplementary Table S1. Comparitive gene duplication content in plasmid pWP5-S18-ESBL-09_1 (NZ_AP022172.1) residing in *E. coli* WP5-S18-ESBL-09**

See supplied Excel file SuppTables.

**Supplementary Table S2. Comparitive gene duplication content in plasmid pUnnamed2 (NZ_CP022035.1) residing in pathogenic *S. enterica* subsp. enterica serovar Onderstepoort SA20060086**

See supplied Excel file SuppTables.

**Supplementary Table S3. Comparitive gene duplication content in plasmid pRHB30-C05_3 (NZ_CP057315.1) residing in *K. pneumoniae* RHB30-C05 isolated from livestock**

See supplied Excel file SuppTables.

**Supplementary Table S4. Comparitive gene duplication antibiotic resistance plasmid pKpN06-CTX (NZ_CP012993.2) residing *K. pneumoniae* KpN06 isolated from human**

See supplied Excel file SuppTables.

**Supplementary Table S5. Comparitive gene duplication content in plasmid pU90-2 (NZ_CP068037.1) residing in *E. coli* U90 hosted by swine with gastroenteritis**

See supplied Excel file SuppTables.

**Supplementary Table S6. Comparitive gene content of a shared segment between plasmid pTEF1 and chromosome of *Enterococcus faecalis* V583 (GCF_000007785.1)**

See supplied Excel file SuppTables.

### Supplementary Figures

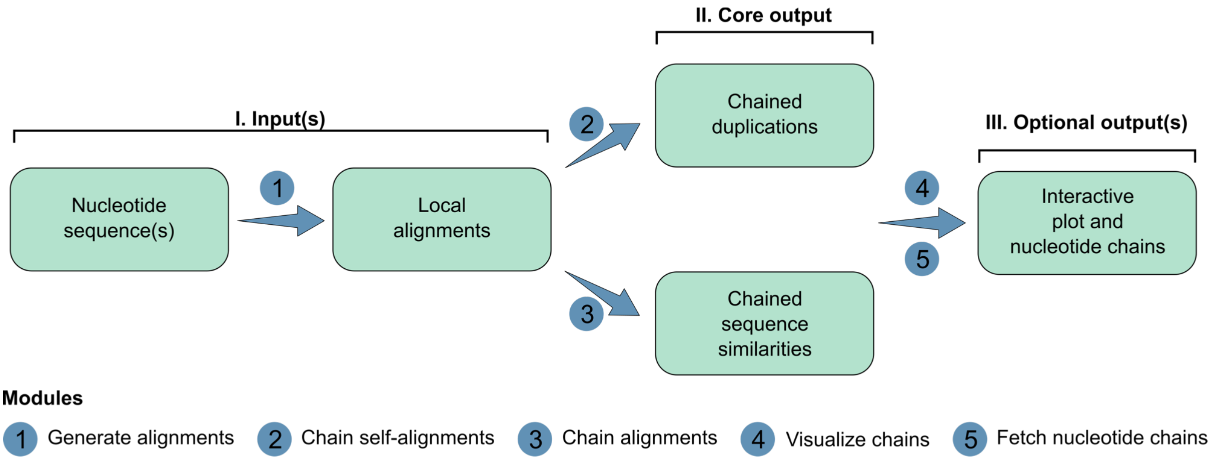

**Figure 1. SegMantX's modules and workflow.** (I) The first module processes nucleotide sequence(s) to compute local alignments, optionally formatting them for further analysis. (II) The core algorithm chains these alignments as duplications or sequence similarities, generating a structured table based on the selected module and purpose. (III) The chained alignments can be explored using the interactive visualization module or extracted as nucleotide sequences for downstream analysis.

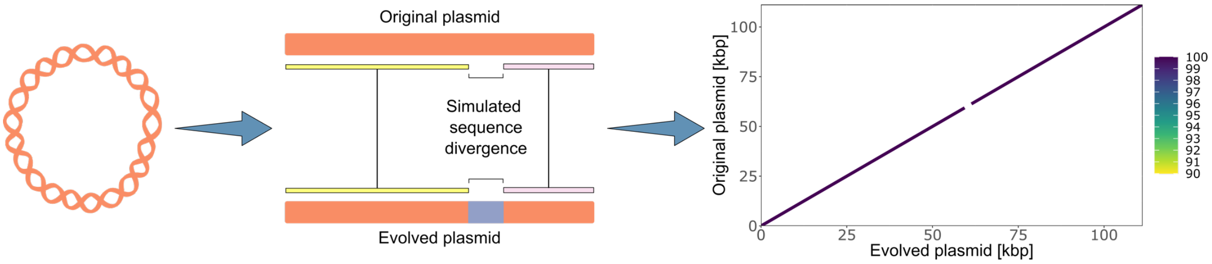

**Figure 2. Graphical illustration of simulated sequence divergence in plasmid genomes.** Individual plasmid sequences were randomly sampled from 1,741 *KES* dataset plasmids lacking significant local alignments in a BLASTn similarity search. Per plasmid sequence single-nucleotide mutations were introduced into randomly selected region, varying in mutated sequence length and mutation frequency. Original and evolved (i.e., mutated) plasmid sequences were then aligned using BLASTn or MMSeqs2 with specified parameters. Alignment continuity between original and evolved sequences was assessed separately for both methods.

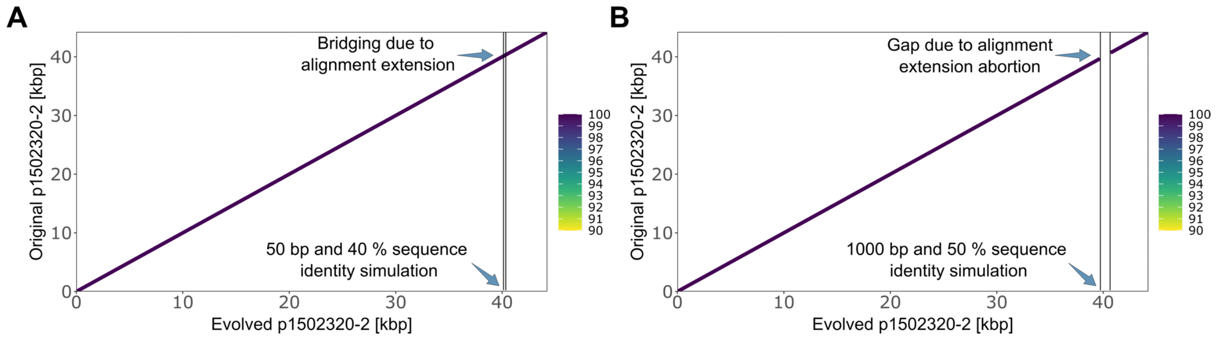

**Figure 3. Example chaining local alignments using simulated plasmid genome sequence data. (A)** A dotplot comparison visualizing BLAST local alignments of plasmid p1502320-2 against its evolved version with a simulated divergence of 40 % (~ 60% sequence identity) within a plasmid genomic region of 50 bp. Note, that aligning the original plasmid to its evolved version using BLAST yield a continuous alignment along the whole plasmid genomes. **(B)** A dotplot comparison visualizing BLAST local alignments of plasmid p1502320-2 against its evolved version with a simulated sequence divergence of 50 % (~ 50% sequence identity) within a plasmid genomic region of 1,000 bp. Note, that aligning the original plasmid to its evolved version using BLAST yield a discontinuous alignment. **(D)** The heatmap depicts simulated gap lengths (x-axis) and gap sequence identity (y-axis). For each pair gap length and gap sequence identity 100 simulations were computed using a plasmid dataset revealing no significant BLAST alignment in a self-alignment search ($n_{plasmids}$= 1,741). The color gradient shows the percentage of simulations that result in fragmented local alignment result from BLAST.

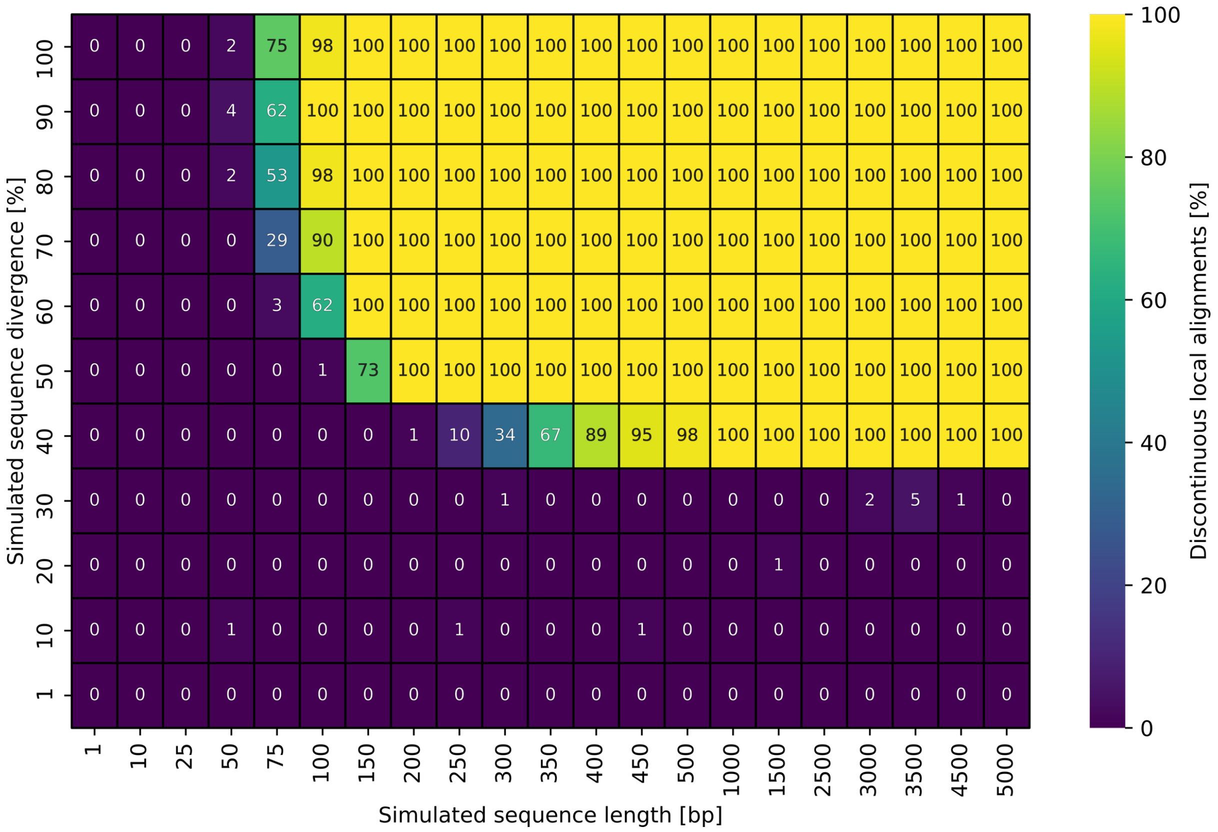

**Figure 4. BLAST alignment continuity results for simulated plasmid genome sequences.** The heatmap illustrates the relationship between simulated gap length (x-axis) and gap sequence identity (y-axis). Each heatmap cell represents 100 simulations using randomly sampled plasmids ($n_{plasmids}$=1,741) that showed no significant BLAST alignment in a self-alignment search. The color gradient indicates the percentage of simulations where BLAST produced fragmented local alignments.

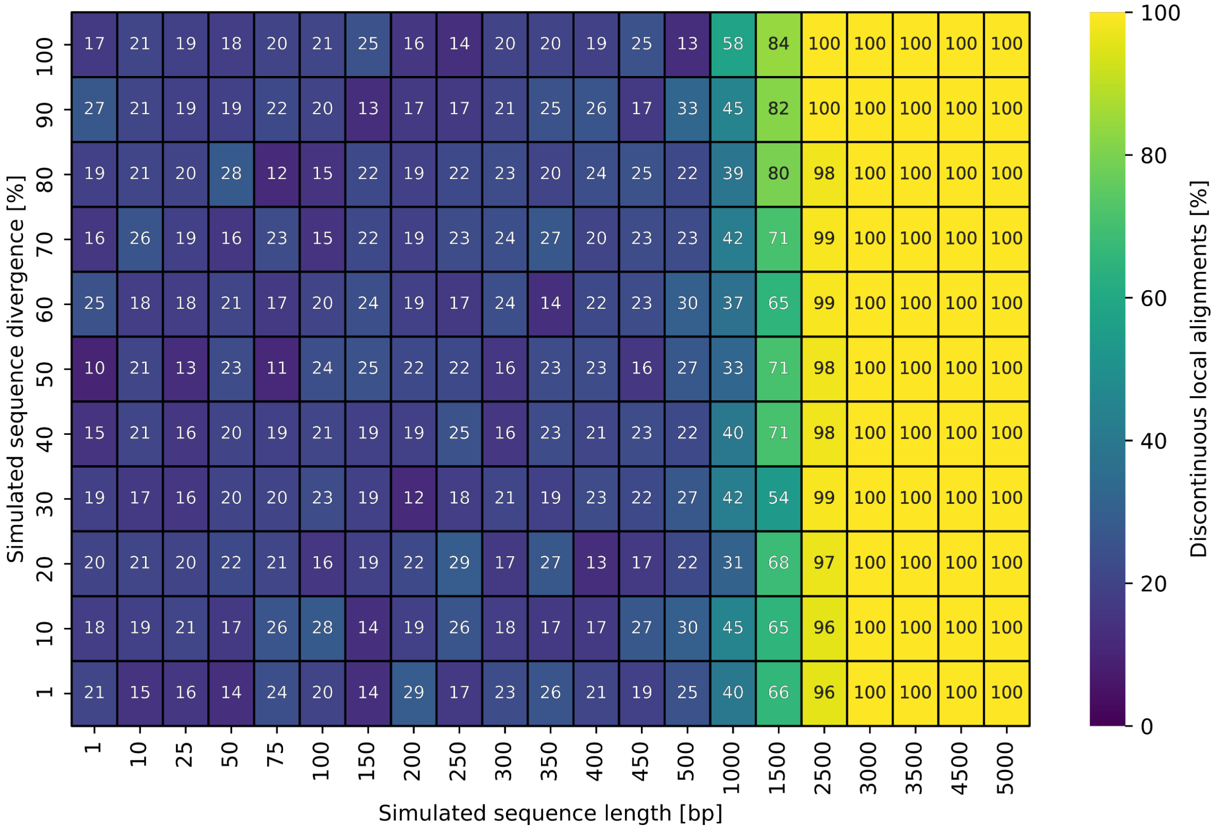

**Figure 5. MMSeqs2 alignment continuity results for simulated plasmid genome sequences.** The heatmap illustrates the relationship between simulated gap length (x-axis) and gap sequence identity (y-axis). Each heatmap cell represents 100 simulations using randomly sampled plasmids ($n_{plasmids}$=1,741) that showed no significant BLAST alignment in a self-alignment search. The color gradient indicates the percentage of simulations where MMSeqs2 produced fragmented local alignments.

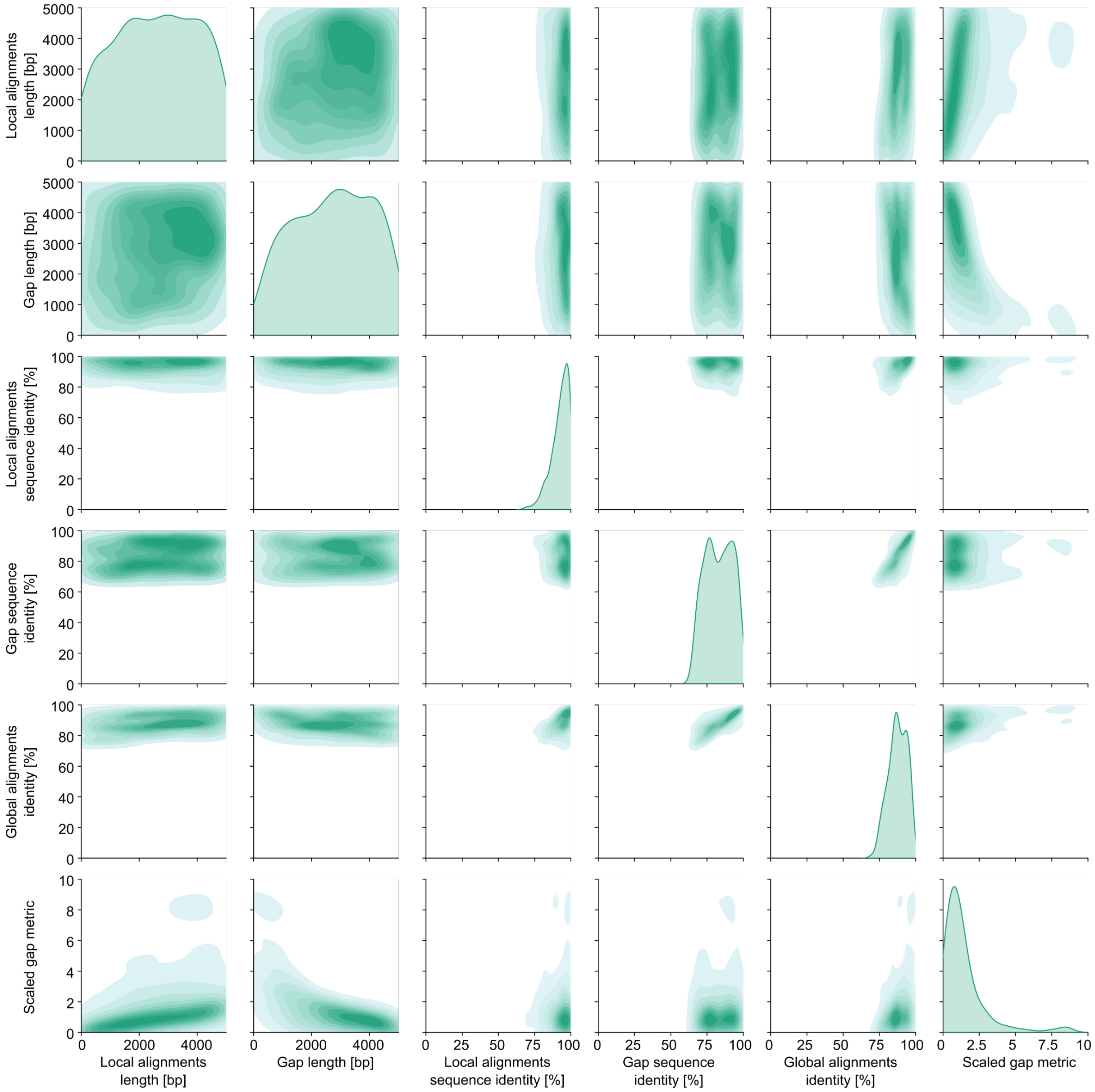

**Figure 6. Pairwise relationship between alignment and gap parameters in simulated plasmid sequences where BLAST successfully produced continuous alignments.** A density plot matrix visualizing the distributions and pairwise dependencies of key alignment and simulation metrics, including local alignment length, gap length, local alignment sequence identity, gap sequence identity, global sequence identity, and a scaled gap metric. Kernel density estimates (KDE) along the diagonal illustrate the correlation structure across these features.

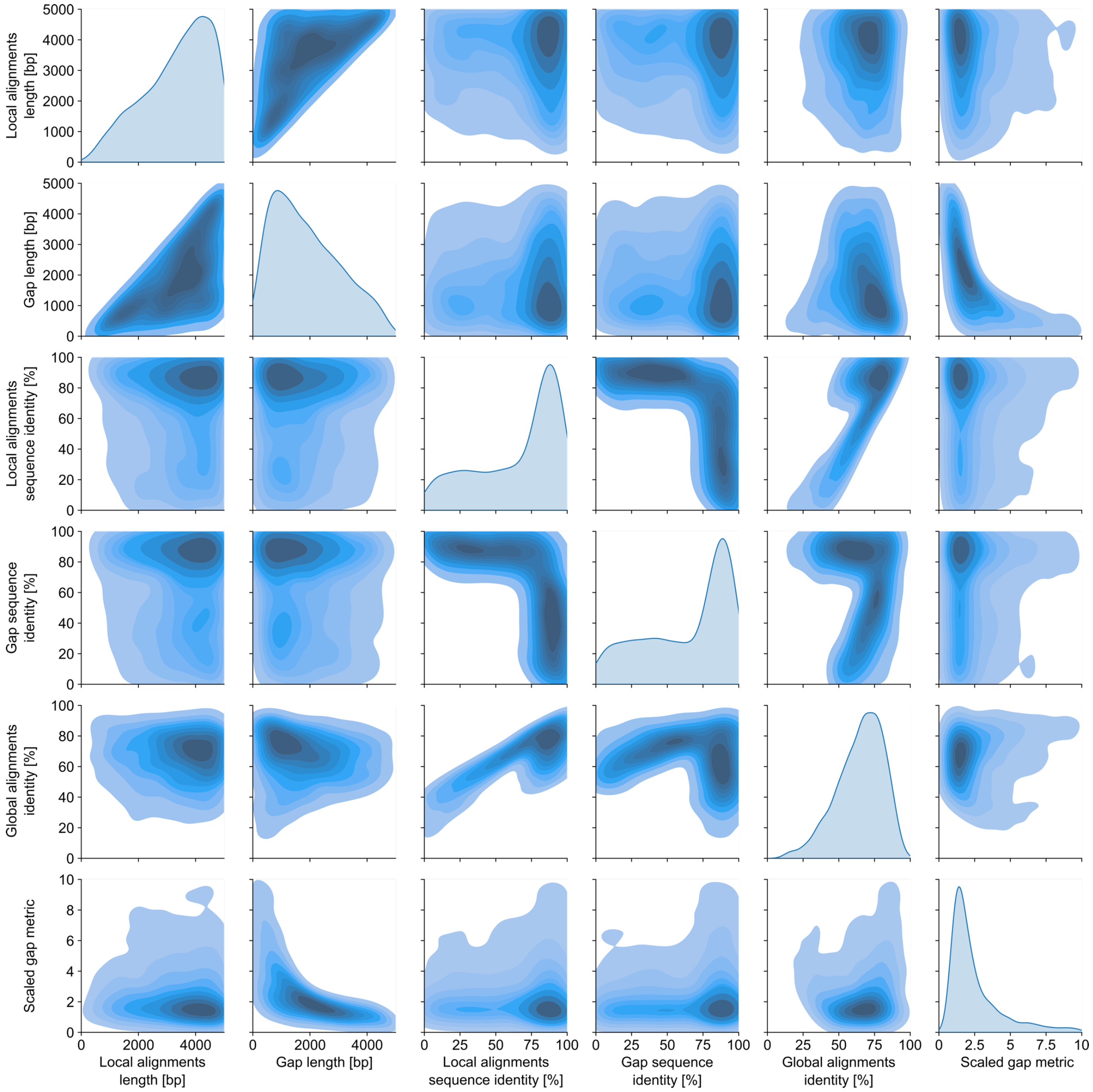

**Figure 7. Pairwise relationship between alignment and gap parameters in simulated plasmid sequences where SegMantX (i.e., local alignment chaining) successfully produced continuous alignments.** A density plot matrix visualizing the distributions and pairwise dependencies of key alignment and simulation metrics, including local alignment length, gap length, local alignment sequence identity, gap sequence identity, global sequence identity, and a scaled gap metric. Kernel density estimates (KDE) along the diagonal illustrate the correlation structure across these features.
